# Virus-host interactions and genetic diversity of Antarctic sea ice bacteriophages

**DOI:** 10.1101/2021.05.28.446129

**Authors:** Tatiana A. Demina, Anne-Mari Luhtanen, Simon Roux, Hanna M. Oksanen

**Author notes:** **Corresponding author**, Molecular and Integrative Biosciences Research Programme, Faculty of Biological and Environmental Sciences, Viikinkaari 9, University of Helsinki, FIN-00014 University of Helsinki, Finland. **Competing interests:**Authors declare no competing financial interests in relation to the work described.

## Abstract

Despite generally appreciated significant roles of microbes in sea ice and polar waters, detailed studies of virus-host systems from such environments have been so far limited by only a few isolates. Here, we investigated infectivity under various conditions, infection cycles, and genetic diversity of Antarctic sea ice bacteriophages: PANV1, PANV2, OANV1, and OANV2. The phages infect common sea ice bacteria belonging to the genera *Paraglaciecola* or *Octadecabacter*. Although the phages are marine and cold-active, replicating at 0-5 °C, they all survived temporal incubations at ≥30 °C and remained infectious without any salts or supplemented only with magnesium, suggesting a robust virion assembly maintaining integrity under a wide range of conditions. Host recognition in the cold proved to be effective, and the release of progeny viruses occurred as a result of cell lysis. The analysis of viral genome sequences showed that nearly half of the gene products of each virus are unique, highlighting that sea ice harbors unexplored virus diversity. Based on predicted genes typical for tailed double-stranded DNA phages, we suggest placing the four studied viruses in the class *Caudoviricetes*. Searching against viral sequences from metagenomic assemblies revealed that related viruses are not restricted to Antarctica, but also found in distant marine environments.

**Importance:** Very little is known about sea ice microbes despite the significant role of sea ice in the global oceans as well as microbial input into biogeochemical cycling. Studies on the sea ice viruses have been typically limited to -omics-based approaches and microscopic examinations of sea ice samples. Up to date, only four cultivable viruses have been isolated from Antarctic sea ice. Our study of these unique isolates advances the understanding of the genetic diversity of viruses in sea ice environments, their interactions with host microbes, and possible links to other biomes. Such information contributes to more accurate future sea ice biogeochemical models.

## Introduction

Sea ice covers a significant area of polar oceans every year, affecting ocean ecology, biogeochemical cycles, and climate (1, 2). Sea ice, especially its liquid brines, is inhabited by various microorganisms, including viruses (3–7) that cope with temperatures below 0° C, rapidly changing salinity, pH, nutrient concentrations, gas fluxes, and varying light conditions (8). Metagenomic studies have revealed high diversity and abundance of viruses in polar aquatic environments (9–11). Virus-like particles concentrations and virus-to-bacterium ratios in sea ice are typically higher than in the surrounding seawater, suggesting active virus production in the ice (12–17). The role of viruses in controlling host abundance is significant in polar environments also due to the lower abundance and diversity of grazers (11, 18). Virus infections in sea ice are not restricted to lytic cycles, but can include lysogenic ones (18, 19) and possibly pseudolysogeny (20). Viruses may also confer properties beneficial to their host survival (21). Low temperature environments in general are suggested to be hot spots of microbial evolution (20).

The studies of sea ice viruses have been typically limited to microscopic examinations and -omics approaches with only a few sea ice virus-host systems isolated, both from the Arctic and the Antarctic (21–24). All the known sea ice bacteriophage isolates display tailed icosahedral virions, except f327, which is filamentous (21–24). Among the sea ice tailed phages, all three types of tails have been observed: long contractile, long non-contractile, and short non-contractile tails, which are characteristics of the myovirus, siphovirus, and podovirus morphotypes, respectively (22–24). In laboratory conditions, the temperature range suitable for the growth of these virus-host systems varies, but the temperature at which the isolated sea ice phages are able to complete a productive infection cycle is typically lower than the maximal growth temperature for their host bacteria (22–24). The isolated sea ice viruses are host-specific or have a narrow host range, and the known hosts are *Shewanella, Flavobacterium, Colwellia, Octadecabacter*, *Glaciecola,* and *Pseudoalteromonas* strains (21–24). The adsorption of two *Shewanella* phages was shown to be fast, as compared to mesophilic phages (25). The Baltic sea ice virus isolates have lytic infection cycles (25), while f327 does not lyse its *Pseudoalteromonas* host, but affects its growth and physiological traits, which might be advantageous to host survival in the natural environment (21). Based on the genome comparisons, the six sequenced Baltic sea ice phage isolates are unrelated or distantly related to each other, except phages 1/4 and 1/40, which are also related to *Vibrio*-specific ICP1-like phages (25). In addition, putative proviral elements related to phage 1/44 were detected in the genomes of *Shewanella* sp. strains (25). The structural proteins of the Baltic sea ice phages recruited translated reads from metagenomic assemblies obtained from various aquatic environments, not being restricted to the Baltic Sea region (25). Taken together, the known sea ice phage isolates suggest that sea ice environments harbor a diversity of phages with complex virus-host interactions, and they are only distantly related to phages from other environments.

Here, we studied four phages isolated from Antarctic sea ice (24) to understand their infectivity under various conditions, genetic diversity, and occurrence in different environments as well as to analyze their infection cycle parameters. Better understanding the role of viruses in sea ice microbial communities would provide valuable information to be included in future sea ice biogeochemical models (26).

## Materials and Methods

### Growth conditions and virus infectivity

Viruses and bacteria (Table 1) were grown aerobically at 4 or 5 °C in Zobell RC medium as previously described (24). The effect of temperature on virus infectivity was tested by incubating virus stocks at 4-55 °C for 1 h. The effects of Na^+^ and Mg^2+^ ions were assessed by diluting virus stocks 1000-fold in SM buffer (50 mM Tris-HCl, pH 7.5, 100 mM NaCl, 8 mM MgSO_4_; (22)), SM buffer lacking either NaCl or MgSO_4_, or both, or lacking MgSO_4_ but supplemented with 10 mM ethylenediaminetetraacetic acid (EDTA) and incubating at 4 °C for 1 and 5 h. The effect of pH was tested similarly using SM buffer either with 50 mM NaH_2_PO_4_ (pH 3, 5, 7.5) or Tris-HCl (pH 7.5, 9). After all incubations, virus infectivity was assessed by plaque assay as previously described (24). Single-factor ANOVA test was used when three or more groups were compared or t-Test (Two-Sample Assuming Equal Variances) when comparing two groups. Groups were considered statistically not different if p>0.05.

**Table 1.** Antarctic sea ice viruses used in this study.

| Virus <sup>a</sup> | Virus morphotype, head diameter | Host strain<sup>a</sup> | Virus genome<sup>b</sup> |  |  |  |  |
| --- | --- | --- | --- | --- | --- | --- | --- |
|  |  |  | Length (bp) | GC% | ORFs | tRNAs | Genbank acc. no |
| Paraglaciecola Antarctic GD virus 1 (PANV1) | Icosahedral head, long contractile tail (myovirus-type), 71±7 nm <sup>a</sup> | *Paraglaciecola* IceBac 372 | 150 766 | 37.7 | 243 | 4 | MW805361 |
| Paraglaciecola Antarctic JLT virus 2 (PANV2) | Icosahedral head, long non-contractile tail (siphovirus-type), 52±8 nm <sup>a</sup> | *Paraglaciecola* IceBac 372 | 35 731 | 41.0 | 73 | 0 | MW805362 |
| Octadecabacter Antarctic BD virus 1 (OANV1) | Icosahedral head, short non-contractile tail (podovirus-type), 68 nm <sup>b</sup> | *Octadecabacter* IceBac 419 | 48 354 | 62.1 | 61 | 0 | MW805363 |
| Octadecabacter Antarctic DB virus 2 (OANV2) | Icosahedral head, short non-contractile tail (podovirus-type), 53±7 nm <sup>a</sup> | *Octadecabacter* IceBac 430 | 39 241 | 53.3 | 76 | 0 | MW805364 |

<sup>a</sup> data from (24);
<sup>b</sup> this study.

### Adsorption and infection cycle

To determine adsorption efficiency and rates, exponentially growing host cultures (OD_550_~0.8) were infected with a multiplicity of infection (MOI) of ~0.001 and incubated aerobically at 4 °C. Samples in which cells were replaced with broth were used as controls. To determine the number of unbound viruses, samples were diluted in 4 °C broth (1:10 or 1:100), cells were removed (Eppendorf table centrifuge, 16 200 g, 5 min, 4 °C), and supernatants were subjected to plaque assay. The percentage of adsorption was calculated from all (particles in broth) and unbound particles (in infected cultures) 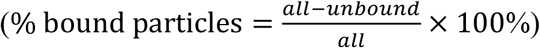. Adsorption rate constant was calculated as 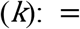 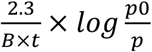, where *B* is cell concentration, *p0* and *p* are free virus concentrations at time point zero and after time period *t*, respectively (27).

For life cycle studies, IceBac 372 (OD_550_~0.8) was infected with PANV1 or PANV2 (MOI 10) and incubated aerobically at 5 °C. Uninfected culture was used as a control. The numbers of infective free viruses in culture supernatants (Eppendorf table centrifuge, 16 200 g, 5 min, 4 °C) were determined by plaque assay. IceBac 419 and IceBac 430 cells (OD_550_~0.8) were collected (Eppendorf table centrifuge, 16 200 g, 5 min, 4 °C) and resuspended in OANV1 or OANV2 virus stocks, respectively, (MOI of ~8) or in broth (uninfected controls). During the growth, the numbers of free viruses and viable cells in supernatant and pellet fractions (Eppendorf table centrifuge, 16 200 g, 5 min, 4 °C) were determined by plaque assay.

### Virus purification and transmission electron microscopy

OANV1 was purified from virus stocks by ammonium sulfate precipitation and rate-zonal ultracentrifugation in sucrose using SM buffer as described (24). Particles were negatively stained with uranyl acetate (2% (w/v), pH 7) or Nano-W (2% (w/v) methylamine tungstate, pH 6.8) prior transmission electron microscopy (Hitachi HT780 microscope, Electron Microscopy Unit, University of Helsinki). Particle size was measured using the ImageJ (28).

### Genome sequencing and annotation

Nucleic acids were extracted from purified viruses by phenol-ether extraction, precipitated by ethanol-NaCl, sequenced using Illumina MiSeq (DNA Sequencing and Genomics core facility, Helsinki Institute of Life Science, University of Helsinki) and assembled with SPAdes version 3.9.0 (29). Sequences are available in the GenBank database (MW805361, PANV1; MW805362, PANV2; MW805363, OANV1; MW805364, OANV2).

Geneious Prime 2021.0.2 (https://www.geneious.com) was used for sequence handling. ORFs were predicted using Glimmer, GeneMarkS (Prokaryotic, v. 3.26), MetaGeneAnnotator (http://metagene.nig.ac.jp/), and FGENESV (http://www.softberry.com/berry.phtml?topic=virus&group=programs&subgroup=gfindv). GC% was calculated with Genomics %G~C Content Calculator (https://www.sciencebuddies.org/science-fair-projects/references/genomics-g-c-content-calculator). Transmembrane helices were predicted using TMHMM Server v. 2.0 (https://services.healthtech.dtu.dk/service.php?TMHMM-2.0). Whole genome comparisons for overall nucleotide identity were done with EMBOSS stretcher (30). Blastn with the whole virus genomes as queries against non-redundant (nr) nucleotide collection (viruses taxid 10239) was used for searching homologous viral genome sequences. Predicted ORFs were assigned with functions based on homology searches with blastx or blastp against nr protein database (thresholds: E-value 0.00001, query cover 30%, identity 30%) (31), blast conserved domains (E-value threshold 0.01) (32), and HHpred within the Toolkit (E-value threshold 0.01) (33) (searches dated 05.2019-02.2021). VIRFAM was used for putative virus classification based on their neck gene module organization (34). tRNA genes were predicted using tRNAscan-SE v. 2.0 (35).

### Metagenomic analyses

Virus sequences were searched against Integrated Microbial Genomes/Virus (IMG/VR) database (36) using the whole genomes as queries and blastn search with the maximum E-value of 0.00001 (search dated 09.12.2020). Sequence similarities were visualized using Circoletto based on Circos (37), with blastn search E-value threshold of 0.00001. For pairwise sequence comparisons, Easyfig v. 2.2.2 was used (38).

The genome sequences of PANV1, PANV2, OANV1, and OANV2 were compared to the IMG/VR spacer databases (both “Isolate” and “Metagenome”) (36) using blastn v2.10.0+ (39) with the following parameters: “-dust no -word_size 7”. Only alignments with 0 or 1 mismatch over the entire length of the CRISPR spacer were considered as potentially informative hits. The corresponding CRISPR spacer were further examined to filter out low-complexity sequences (e.g., short predicted spacers including repeat sequences).

## Results

### Antarctic sea ice phage isolates PANV1, PANV2, OANV1, and OANV2 tolerate elevated temperatures and lowered salinity

To assess virus infectivity at different temperatures, viruses were incubated at 4 to 55 °C, using 4 °C as a reference (100% infectivity) (Fig 1). All four viruses stayed fully infectious when temporally exposed to the temperatures up to 30-45 °C. No statistically significant difference was observed in PANV1 infectivity at 4 and 35 °C, whereas the titer dropped to ~2 % at 40 °C, and only ~0.001% of particles were infective at 45 °C. For PANV2, 45 °C temperature had no statistically significant effect on the infectivity, but a sharp titer drop to ~0.03% was observed at 50 °C. OANV1 preserved the titer at 25 °C, while at 30 and 35 °C, the infectivity was ~28% and ~7.5%, respectively. For OANV2, no statistically significant difference in titers at 4 °C and 40 °C was observed, but only ~4% of particles were infective after incubating at 45 °C. Virus titers decreased one-tenth or more at 40, 50, 35, and 45 °C for PANV1, PANV2, OANV1, and OANV2, respectively. The titers were under the detection limit (<1×10^3^ PFU/ml), at 50, 55, 40 and 50 °C for PANV1, PANV2, OANV1, and OANV2, respectively (Fig 1).

**Figure 1.**
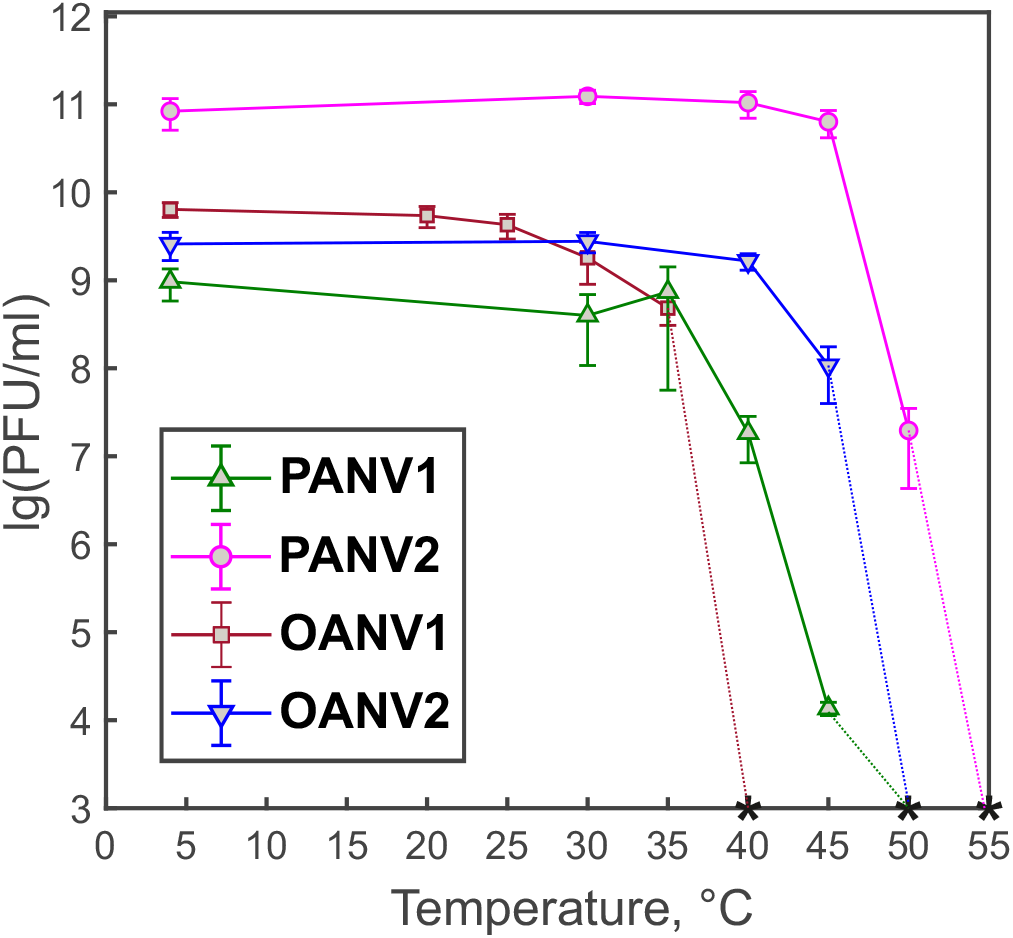
Virus infectivity after 1 h exposure to different temperatures. Bars represent means of at least three independent replicates with standard error of the mean. Asterisks indicate that the titers are under the detection limit (<1×10^3^ PFU/ml).

All four viruses studied here remained infectious without NaCl (Fig. 2 a–d, buffer 2). Moreover, PANV1 and PANV2 infectivity was preserved in the absence of both NaCl and Mg^2+^, even if residual Mg^2+^ ions were removed by the chelating agent, EDTA (Fig. 2 a, b, buffers 3-5). When only MgSO_4_ was excluded from SM buffer, OANV1 titer dropped by 4-5 orders of magnitude, and OANV2 titer dropped to one-tenth (Fig. 2 c, d, buffer 3). If removing all residual Mg^2+^ ions using EDTA (Fig. 2 c, d, buffer 4), OANV1 titer was under the detection limit (<1×10^4^ PFU/ml), and OANV2 titer dropped 100-fold. When both NaCl and MgSO_4_ were removed (Fig. 2 c, d, buffer 5), OANV1 and OANV2 titers were <1×10^4^ PFU/ml.

**Figure 2.**
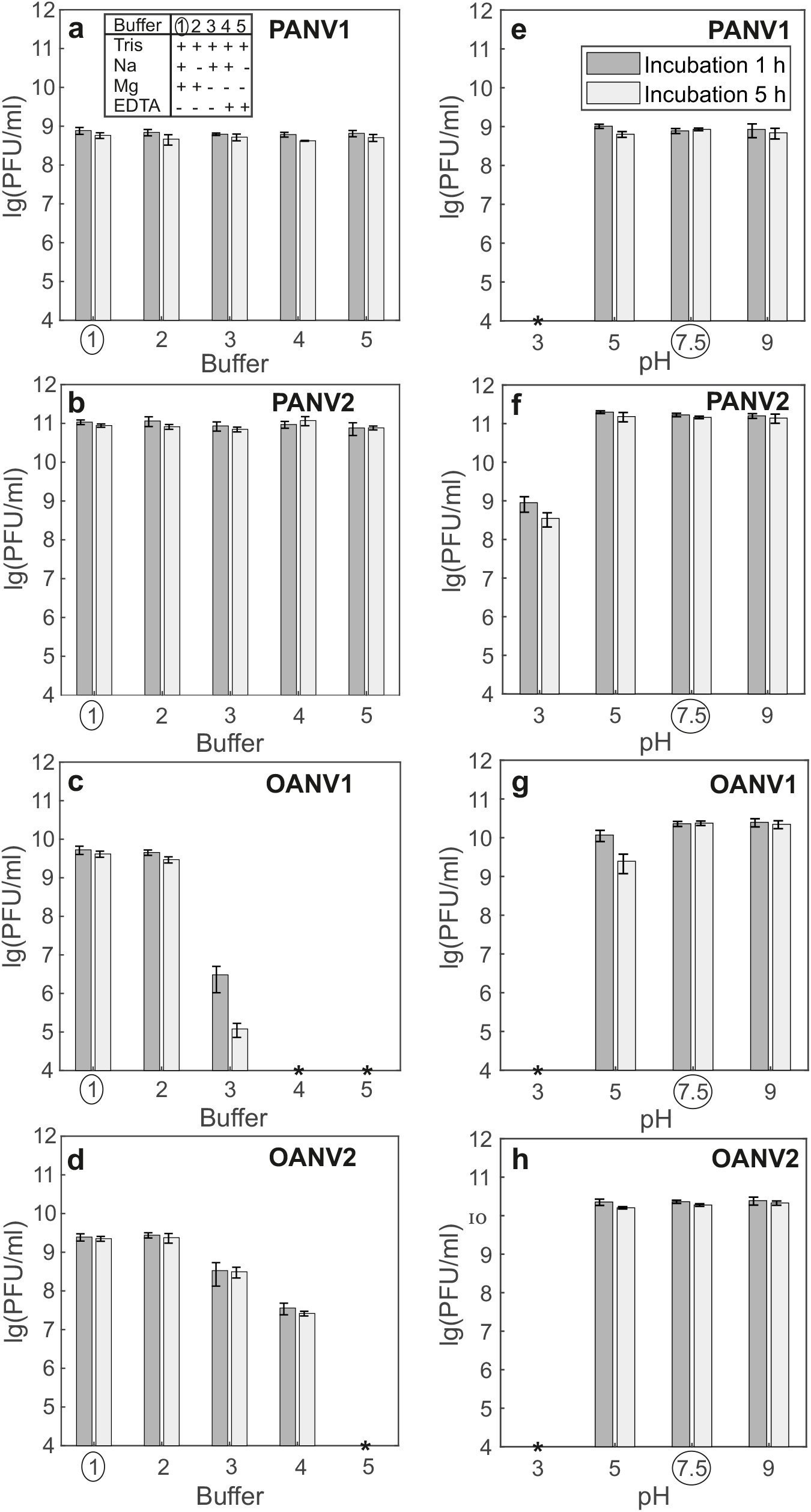
Virus infectivity in buffers with different ion composition and pH. Virus stocks were diluted either in SM buffer (circle in all panels), (a-d) modified SM buffers where some components were omitted (key in a), or (e-h) modified SM buffers of different pH. The incubation lasted 1 h (dark grey) and 5 h (light grey). Bars represent means of at least three independent replicates with standard error of the mean. Asterisk indicates that the titer is under the detection limit (<1×10^4^ PFU/ml).

No statistically significant difference in virus titers was observed when virus stocks were incubated in SM buffer pH 7.5 either with Tris-HCl or NaH_2_PO_4_ (Fig. S1). For all four viruses, pH 9 had no significant effect on the infectivity (Fig. 2 e–h). PANV1, PANV2, and OANV2 were also stable at pH 5. OANV1 preserved the titer at pH 5 after 1 h incubation, but the titer dropped 10-fold after 5 h. At pH 3, the titers dropped significantly for all four viruses (PANV2 <1%; others <1×10^4^ PFU/ml).

### Antarctic sea ice phages PANV1, PANV2, OANV1, and OANV2 adsorb to their hosts effectively

PANV1, PANV2, OANV1, and OANV2 effectively adsorbed to their hosts, achieving at least ~50% binding efficiency within 6 h at 4 °C (Fig. 3). During the experimental set-up, the viruses were not inactivated, since no decrease in virus plaque numbers was observed in virus control samples. PANV2 and OANV2 showed the fastest and the most efficient adsorption, having ~80% particles adsorbed by 30 min and 1 h post infection (p.i.), respectively, and reaching ~100% binding later (Fig. 3). Adsorption rate constants *k* calculated for the first 30 min p.i. (n=3) were 3.9×10^−9^ and 9.0×10^−12^ ml/min for PANV2 and OANV2, respectively. PANV1 and OANV1 adsorbed with the rates of 5.4×10^−10^ and 4.6×10^−13^ ml/min, respectively, reaching ~70% adsorption by 12 h p.i.

**Figure 3.**
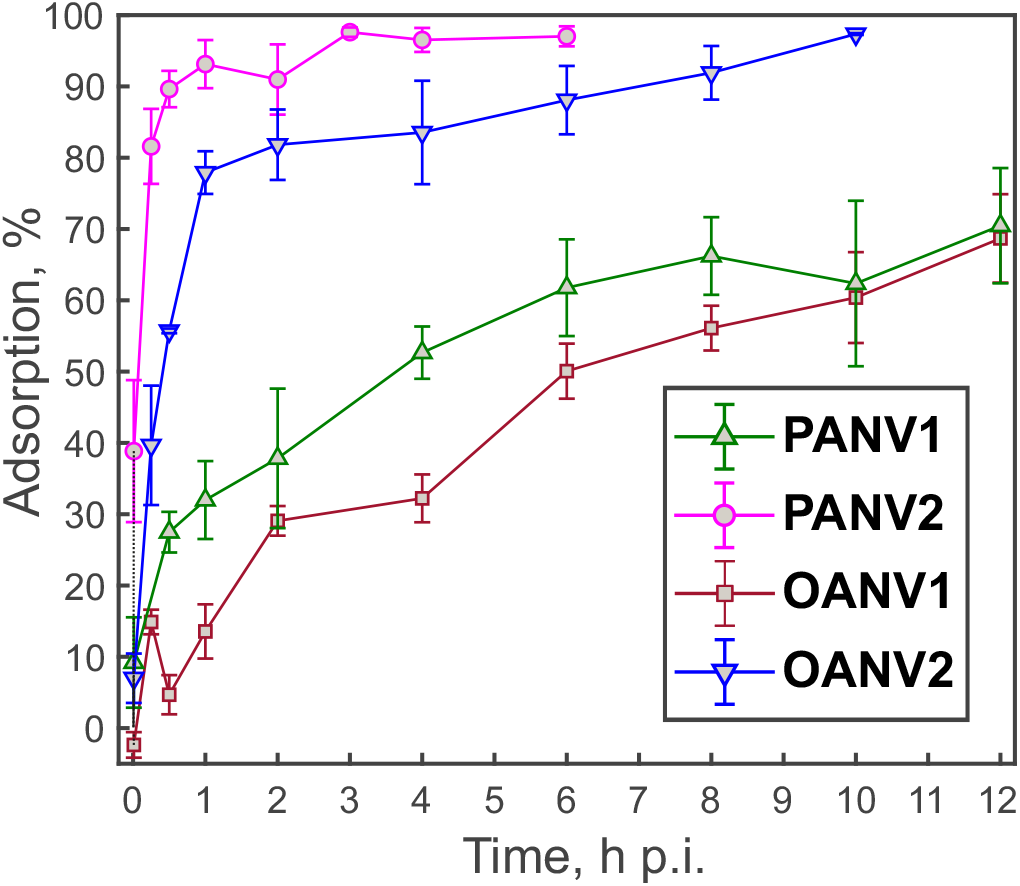
Adsorption efficiency shown as percentage of bound viruses at 4 °C. Means of at least three independent replicates are presented with standard error of the mean.

### Infection cycles of PANV1, PANV2, OANV1, and OANV2 phages result in cell lysis

Bacterial strains *Paraglaciecola* IceBac 372 and *Octadecabacter* IceBac 419 and 430 grew to early stationary stage in 4-7 days (from OD_550_=0.2 to 1.4-1.6), showing typical growth curves of a bacterial culture. The cultures were infected at the logarithmic growth phase (IceBac 372: OD_550_=0.8, ~2×10^7^ CFU/ml; IceBac 419 and IceBac 430: OD_550_=0.6, ~3×10^9^ CFU/ml) using a MOI of 8-10 to analyze one-step growth of the phages (Fig. 4). Uninfected cultures reached OD_550_=1.6-1.7 by 124 h p.i. (Fig. 4). The turbidities of cultures infected with PANV1, PANV2, or OANV2 started to decrease at 12-20 h p.i, and eventually dropped to 0.2-0.4, indicating cell lysis (Fig. 4 a, b, d). The optical density of OANV1-infected culture stayed at the same level as at the time of infection (Fig. 4c). For PANV1 and PANV2, an increase in the numbers of free viruses was detected at 24 h p.i., suggesting progeny virus production. The lysate titers were ~1.6×10^10^ and ~2.4×10^11^ PFU/ml for PANV1 and PANV2, respectively. In OANV1 and OANV2 infections, the increase of free viruses could not be detected with the methods used here. However, for OANV1, the number of viable *Octadecabacter* IceBac 419 cells at the time of infection (~1.1×10^9^ CFU/ml) reduced almost two orders of magnitude as a result of the virus addition (~2.3×10^7^ CFU/ml) by 124 h p.i., while the number of viable cells in the uninfected culture of IceBac 419 was growing (~3.3×10^9^ CFU/ml by 124 h). This can be interpreted as lysis caused by the virus infection. Similarly, the numbers of viable cells in the OANV2-infected IceBac 430 culture dropped noticeably (from ~1.4×10^9^ at 0 h p.i. to ~5×10^6^ CFU/ml at 124 h p.i.) compared to the uninfected culture (~3.5×10^9^ CFU/ml at 124 h p.i.).

**Figure 4.**
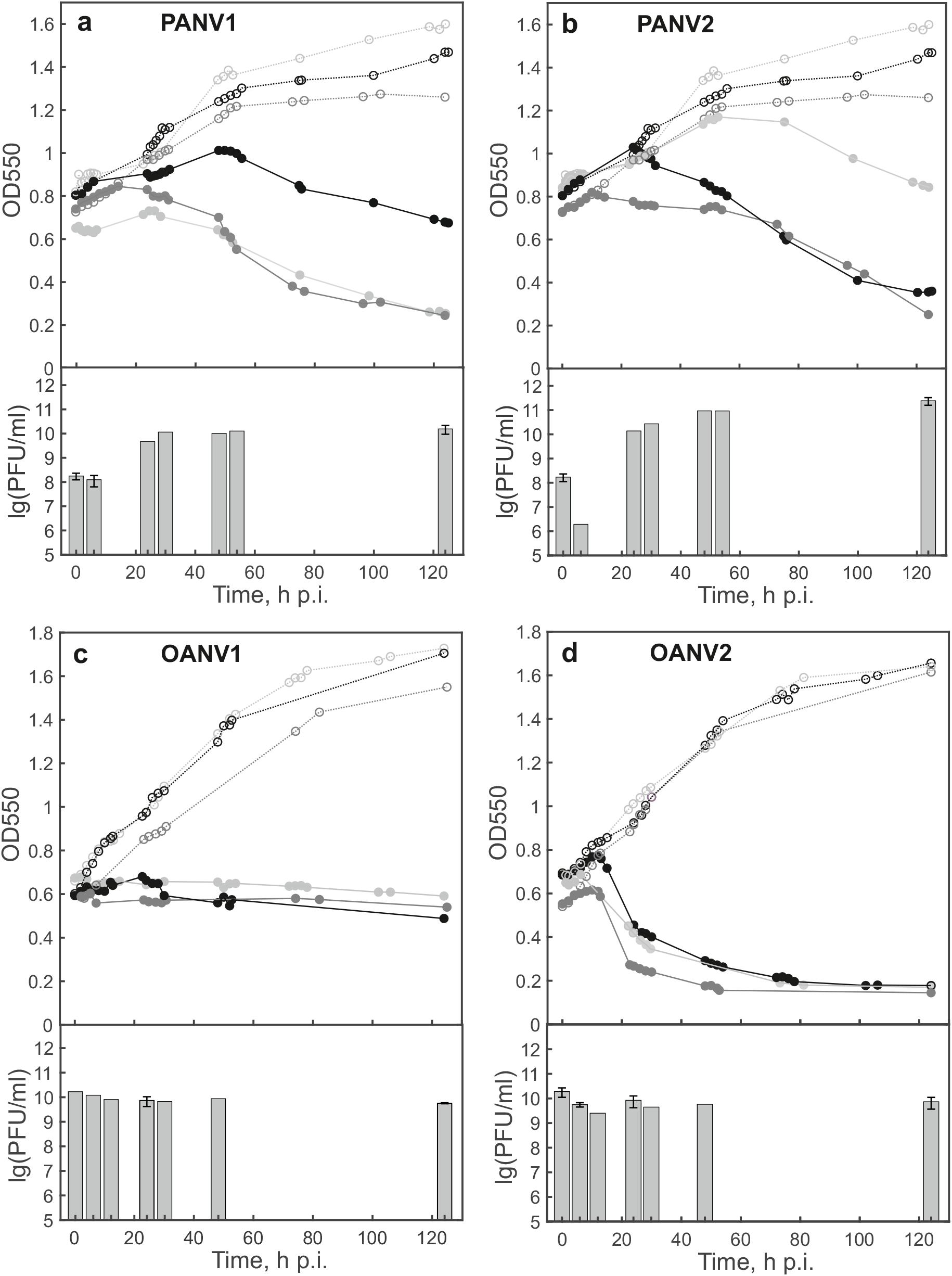
One-step growth curves of (a) PANV1 and (b) PANV2 in *Paraglaciecola* IceBac 372, (c) OANV1 in *Octadecabacter* IceBac 419, and (d) OANV2 in *Octadecabacter* IceBac 430. Growth curves of uninfected (open circles, dashed lines) and infected (closed circles, solid lines) cultures from three independent repeats are shown in the upper panels. Curves with same shades of grey represent the same repeat. The numbers of free viruses are shown as bars in lower panels: means with standard error of the mean where appropriate (n=3), otherwise means of n=2.

### The four Antarctic sea ice phage isolates have largely unique genomes

PANV1, PANV2, OANV1, and OANV2 genomes are dsDNA molecules ranging from ~36 to ~151 kb, with GC content of 38-62% and 61-243 predicted protein-coding ORFs (Table 1, Fig. 5a, Tables S1-4). Of the phages studied here, only PANV1 contains predicted tRNA genes (Table 1). All four virus genomes include ORFs encoding small and large terminase subunits, which are hallmark genes for tailed dsDNA bacteriophages that package their genomes into a preformed procapsid (40). ORF numbering in four genome sequences studied here was started with the ORF for the small terminase subunit (ORF1).

**Figure 5.**
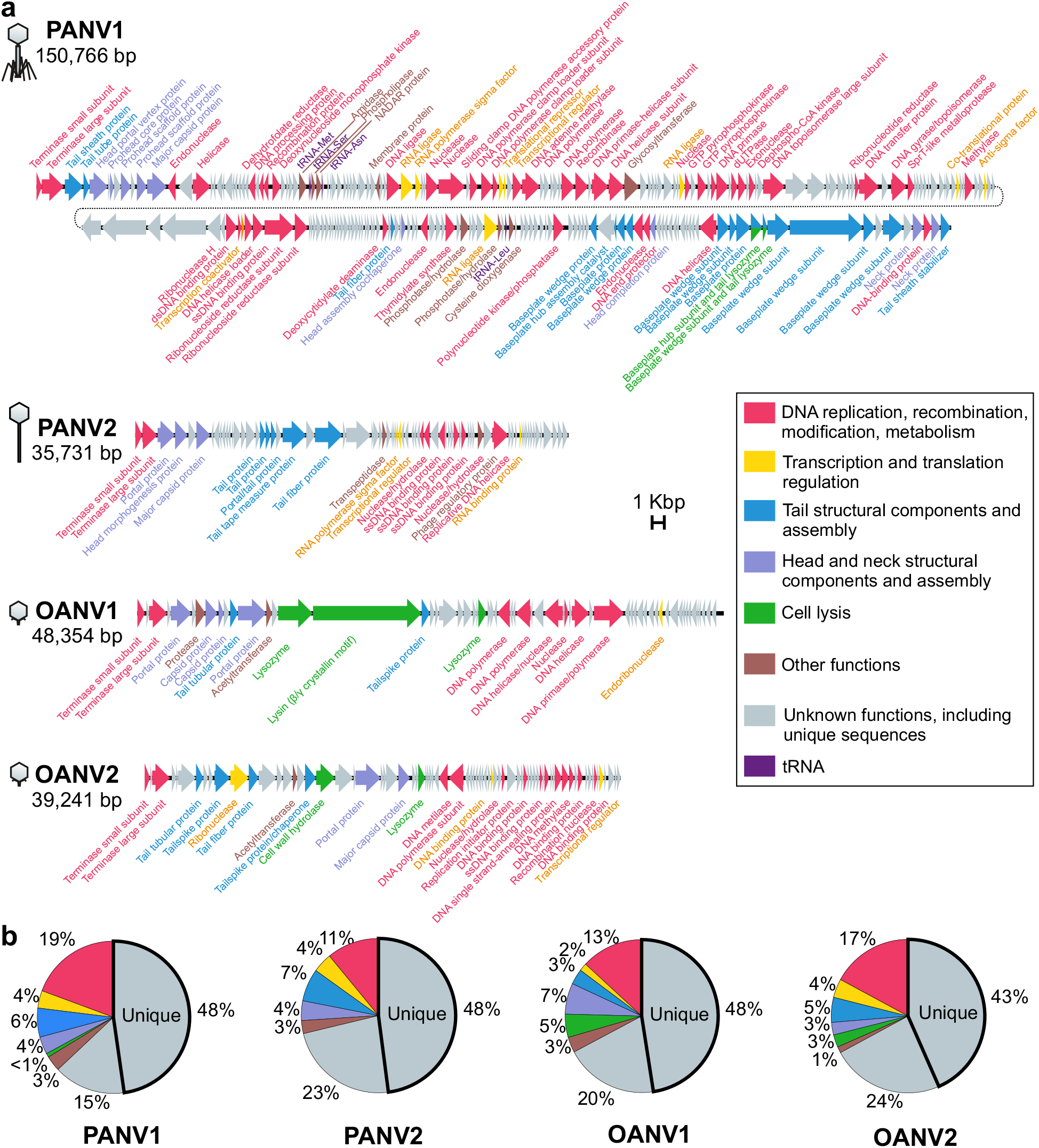
(a) Genomes of PANV1, PANV2, OANV1, and OANV2. ORF colors refer to the assigned functional categories (inset, same for (a) and (b)). Virus morphotypes are indicated schematically on the left. (b) Distribution of functional categories assigned to the protein-coding ORFs of PANV1, PANV2, OANV1, and OANV2. The portions of unique ORF products not having homologs in the NCBI nr protein database are outlined with bold.

The majority of predicted ORF products (63-71%) of the four virus genomes could not be assigned with any functions, including unique sequences (43-48%) that had no homologues in the NCBI nr protein database (Fig. 5b). Surprisingly, the ratio between unique ORF products and those that had some homologous sequences in the database was about the same for all four viruses, regardless of the genome length (Fig. 5b).

About 29-37% of the studied sequences had homologs in the database, allowing us to predict their functions. Blast matches revealed mosaic similarities to the sequences of other phages, as well as moderate and psychrophilic bacteria. The following functional categories were assigned to the gene products (gp): (i) proteins involved in DNA replication, recombination, modification, and metabolism; (ii) transcription and translation regulation proteins; (iii) virion and tail structural components; (iv) cell lysis; and (v) other functions (Tables S1-S4, Fig. 5). Some putative proteins could be classified to more than one category.

PANV2 gp55 has significant blast hits to phage regulatory Rha proteins, encoded by temperate phages (41, 42). Unlike the other three viruses, no lysis genes were predicted in PANV2. In OANV1, gp15 and gp16 are putatively lysis-related proteins. Noticeably, gp16 is the longest predicted OANV1 protein (2 986 residues) and it is similar to *Bordetella* phage BPP-1 bbp10, which is a lysin containing beta/gamma crystalline motif (blastp search 19.02.21: 50% cover, 30% identity, E-value: 3e-128). In PANV1, gp232 and gp233 were annotated as baseplate subunits having lysozyme activity, being similar to the gene products in T4-like phages (T4 gp5 and gp 25, respectively) (43, 44). In OANV2, gp19 is presumably a cell wall hydrolyze and gp29 is a lysozyme.

Along with the categories typical for dsDNA phage genomes, additional functions were also predicted. PANV1 gp36 was predicted to be a phospholipase (HHpred search: hit 1LWB_A, probability 98.8, E-value 2e-8, 19.2.2021), thus possibly involved in lipid metabolism. PANV1 gp37 putatively belongs to the NADAR (NAD and ADP-ribose) superfamily, having a hydrolyse activity and taking part in carbohydrate derivative metabolic processes (HHpred search: hit 2B3W_A, probability 100, E-value 7.1e-34, 19.02.2021). PANV1 gp51 had matches to mechanosensitive channel proteins, involved in transmembrane transport (e.g., Hhpred hit to 6RLD_D, probability 99.7, E-value 7.2e-17, 19.02.2021). However, a transmembrane helix was predicted in this protein with only ~0.6 posterior probability by TMHMM v 2.0. In PANV2, gp30 is a putative transpeptidase, involved in peptidoglycan cross-linking (HHpred, 4LPQ_A, probability 96.6, E-value 0.01, 19.02.2021).

Based on the Virfam analysis of the neck module and part of the head and tail proteins (34), PANV1 was assigned to the category of “*Myoviridae* of Type 2”, adopting the structural organization of the myophage T4 neck. PANV2 was assigned to “*Siphoviridae* of Type 1 cluster 5”, adopting the structural organization of the siphophage SPP1 neck. Both OANV1 and OANV2 were predicted to belong to “*Podoviridae* of Type 3”, adopting the structural organization of the podophage P22 neck. (34). The Virfam-based classification is consistent with the tail morphology determined by transmission electron microscopy previously for PANV1, PANV2, and OANV2 (24) and here for OANV1 (see below).

### OANV1 is a podovirus

Since the sequence information indicated that OANV1 is not a siphovirus, we re-analyzed the OANV1 virus morphology. Transmission electron micrographs of purified OANV1 particles displayed a podovirus-like morphotype: tailed virions with icosahedral heads (diameter ~68 nm, n=44) and short non-contractile tails (length ~11 nm, n=25) (Figs. S2a and S2b). Electron micrographs were taken from two independently purified virus samples (specific infectivity 6.1×10^13^ and 3.7×10^13^ PFU/mg of protein) using two negative stains, all demonstrating consistent particle morphology. The protein patterns of the purified OANV1 particle samples were identical to each other (Fig. S2c) and similar to that previously reported (24). OANV1 was initially identified as a siphovirus most probably due to some error during the sample preparation and imaging (24).

### Viruses related to the known Antarctic sea ice phage isolates are found in Antarctica and in other distant marine environments

The overall nucleotide identity between four genomes is the lowest (~24%) between PANV1 and PANV2 and the highest (~52%) between OANV1 and OANV2. When the complete sequences were used as queries in blastn searches (somewhat similar sequences option, dated 19.02.2021) against all virus sequences (NCBI Taxonomy ID10239) in the nr nucleotide collection, typically only less than 3% of the whole genome sequences could be aligned with other virus sequences available, highlighting the overall uniqueness of these Antarctic sea ice virus isolates. One exception was the match of the OANV2 genome to the uncultured *Caudovirales* phage genome assembly (Genbank accession number LR798304), obtained from metagenomes from the Římov reservoir (freshwater man-made pond), Czech Republic (45). The overall identity between the genomes was 52% (Fig. S3a).

In contrast, a similar blastn search with PANV1, PANV2, OANV1, and OANV2 whole genomes as queries against the IMG/VR (36) database resulted in many hits to sequences obtained from various locations and environments. We have given a priority to the search based on the whole genome sequences rather than separate ORFs to ensure more specific hits and the possibility to easily select matches with several regions of similarity. Both regions containing ORFs with assigned functions and unknown ones recruited hits. Most blastn hits covered relatively short regions (38-3870 nt), hence, to find similar viral genomes rather than separate ORFs, scaffolds with at least three regions of similarity were selected for further analysis (Table S5). While PANV1 had no such related scaffolds, the other three virus genome sequences recruited several scaffolds originating from Antarctica and other environments (Fig. 6). Notably, some of the selected scaffolds represented identical parts of each other and originated from the same project and sampling location, so likely represented sequencing of the same virus across multiple samples (the duplicates excluded in Fig. 6).

**Figure 6.**
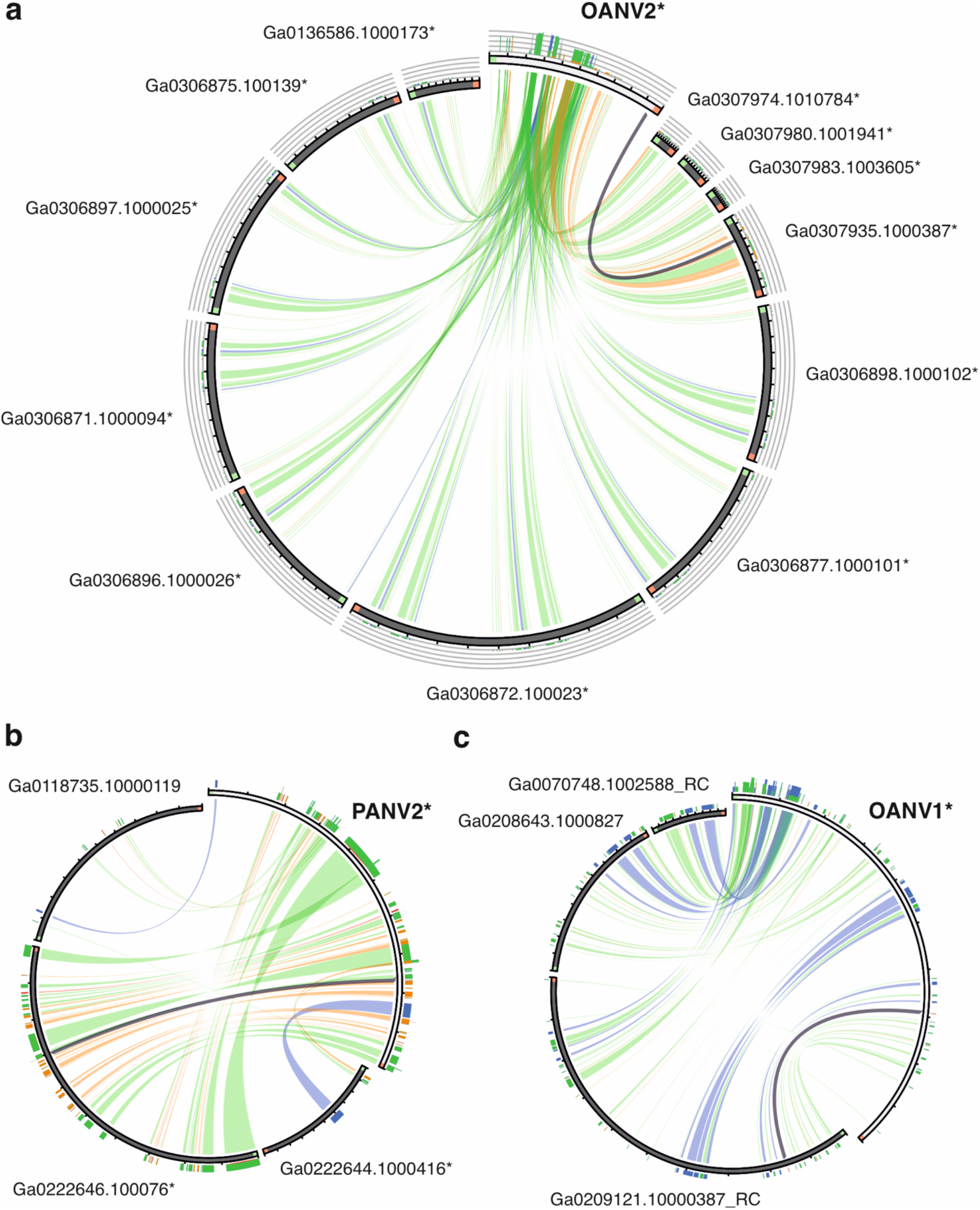
Visualizing sequence similarities between (a) OANV2, (b) PANV2, and (c) OANV1 and corresponding metagenome-derived scaffolds found in IMG/VR database (see Table S5). The figure was generated with Circoletto, colouring ribbons by % identity with absolute colouring: blue<=80, green<=90, orange<=95, red>95. Minimal and maximal identity intervals are (a) 77.9-100.0%, (b) 79.2-100.0%, and (c) 78.3-97.0%. Orientation of the sequences is clockwise, in addition, sequence starts are marked with green and ends with red. Some sequences are presented as reverse complements (RC). Sequences originating from Antarctica are marked with asterisk.

PANV2 was similar to scaffolds assembled from saline water microbial communities from Ace Lake, Antarctica (46), and one scaffold from marine sediment microbial communities from methane seeps sampled in Hudson Canyon, US Atlantic Margin. OANV1 was related to scaffolds from oil polluted marine microbial communities from Coal Oil Point, Santa Barbara, California, USA (47, 48), and aqueous microbial communities from the Delaware River and Bay (49), but no samples from Antarctica (Fig. 6). On the contrary, OANV2 was similar only to scaffolds originating from saline lake microbial communities from various locations in Antarctica: Organic lake, Club lake, Deep lake, and saline lake on Rauer Islands (46, 50, 51). Similarity regions are distributed along the whole virus genomes and include ORFs assigned with different functions, including those encoding capsid structural proteins (Figs. S3b, S4, S5). The overall nucleotide identity between the query genomes and scaffolds retrieved from the database was ~40% for those sequences that have similar length (Table S5).

A blast search of PANV1, PANV2, OANV1, and OANV2 genomes against the IMG/VR spacer databases did not yield any information about potential additional hosts of these viruses: no reliable hits were detected to CRISPR spacers from isolate genome, and the hits to metagenome-derived spacers were on short metagenome contigs that could not be taxonomically affiliated.

## Discussion

### The effect of elevated temperature, low ionic strength, and acidic/alkaline conditions on viral infectivity of the known sea ice phage isolates

The sea ice phages studied here were able to cope with temporal exposures to the temperatures of at least 30 °C or even 50 °C (Fig. 1), which is much higher than their growth temperature (0-5 °C) (24). Similar physical stability of virions has been observed in the tailed dsDNA phages isolated from the Baltic sea ice on *Shewanella* or *Flavobacterium* (23, 24) and cold-active phages isolated from Napahai wetland and Mingyong glacier, China (52–56). In comparison, the cold-active tailed dsDNA phage 9A of *Colwellia psychrerythraea* from the Arctic nepheloid layer is rapidly inactivated at 25-33 °C with no effect of salinity or clay particles on thermal lability (57). The sea ice viruses studied here demonstrated no infectivity losses in the absence of NaCl, which is the main salt in marine water. Moreover, PANV1 and PANV2 infectivity was not affected by the absence of Mg ions either. Maintaining of infectivity in the absence of salts as well as in a wide pH range may be beneficial for the viral survival in sea ice brine channels, where salinity and pH may change across the brine network (6, 8). Some bacteriophages isolated from solar salterns, where salinity may considerably change following evaporation and rainfall events, also have wide salinity tolerance ranges (58, 59), while others are more sensitive to lowered salinity (60). Similarly to the observed lack of typical patterns in virion stability of cold-active phages, different patterns in the variation of infectivity have been observed in bacteriophages in general, not depending on virus morphotype or taxonomic assignments (61). Virus thermal inactivation may be caused by the release of genetic material from the capsid, as well as DNA and protein denaturation (62). Bacteriophages may aggregate at pH levels lower than their isoelectric point (63) or in lowered ionic strength conditions (64). The stability of PANV2 under different conditions shown here makes it an attractive model for future studies of molecular adaptation and virion architecture of viruses residing in sea ice.

### Host interactions of the Antarctic sea ice virus isolates

Viruses isolated from Arctic, Baltic and Antarctic sea ice so far seem to be very host-specific and have different temperature limits for successful host infection, as tested in laboratory conditions (22–24). No new potential hosts could be assigned to the viruses studied here using the IMG/VR spacers databases, which likely reflects the specificity of sea ice phages and the limited representation of Antarctic microbes in the current genome and CRISPR spacer data. The relatively fast adsorption of PANV2 (90% in 30 min) and slower adsorption of PANV1 (maximally ~70% in 12 h) resulted in non-synchronized lysis and active virus production in *Paraglaciecola* observed at 24 h p.i. Putative tail fiber proteins identified in PANV1 may have roles in the attachment to host cells. The similarity of the PANV1 gp232 and gp233 to lysozymes in T4-like phages suggests similar lysis mechanisms (43, 44), whereas siphovirus PANV2 may use some unknown lysis mechanisms, since no hits to known lysis-related proteins were identified, or its efficient lysis might be dependent on another resident phage (65). Lysogeny seems to be prevalent in polar regions (19, 66, 67). The similarity of PANV2 gp55 to Rha regulatory proteins found in lambdoid phages and bacterial prophage regions (41, 42) suggests that PANV2 may establish lysogenic infection cycle. Switches between infection modes can be triggered by environmental factors as well as the physiology of host cells (68).

For *Octadecabacter* podoviruses, OANV2 adsorption was faster and more efficient than that of OANV1 to their respective host strains. Putative tail spike proteins, possibly involved in the attachment to host cells, were found both in OANV1 and OANV2. OANV2 infection resulted in a sharp cell density decrease at 12 h p.i., while no decrease in the optical density was observed for OANV1 infected culture. Nonetheless, the number of viable cells decreased during both infections, suggesting cell lysis, which is also supported by the presence of putative lysis-related genes in OANV1 and OANV2 genomes.

Adsorption rate constants of four Antarctic sea ice viruses varied noticeably from the fast binder PANV2 to the slow binder OANV1 with PANV1 adsorption rate of 5.4×10^−10^ ml/min being the most similar to that observed previously in *Shewanella* phages isolated from Baltic sea ice, 1/4 and 3/49, at 4 °C (25). Infection cycles of the phages 1/4 and 3/49 were, however, considerably faster (measured at 15 °C) (25). Cold-active tailed bacteriophages from Napahai wetland and Mingyong glacier, China, have optimal plaque formation at 15-20 °C, rapid and efficient adsorption, short latent period, as well as fast and complete lysis (52–56). Thus, Antarctic sea ice phages studied here have effective, but relatively slow infections, which may be due to slow host growth rates and preferred low temperatures (24). Similarly, slow infections with long latent periods have been observed for the *Pseudoalteromonas* phage from the North Water, Arctic, (15 h at 0 °C) (69) and *Pseudomonas putrefaciens* phage 27 from Boston harbour water (8.5 and 14 h at 2 °C on strains P10 and P19X, respectively) (70). In case of cold-active bacteriophage 9A infection in *Colwellia psychrerythraea* 34H, the latent period was shown to differ depending on the growth temperatures, ranging from a few hours at 8 °C to several days at temperatures below zero (71). Phage adsorption rates and latent period length may also depend on the host pre-incubation temperature (70, 72).

### Genetic diversity of Antarctic sea ice virus isolates and possible links to other biomes

Viral diversity seems to be unique for Antarctica, but generally lower than that in lower-latitude marine systems (73). Overall, Antarctic phages studied here are genetically diverse and not closely related one to another or to any other sequenced sea ice phage isolates (22, 25, 74). The detected genetic similarities are rather mosaic and are not restricted to cold-active microorganisms. The identified functional categories are typical for tailed dsDNA phages, and we suggest placing four studied viruses in the class *Caudoviricetes*. A few PANV1 proteins may take part in lipid and carbohydrate metabolism and membrane transport, thus possibly being auxiliary metabolic genes, which is commonly seen in marine phages (75). High percentage of unique sequences in the Antarctic sea ice virus isolates emphasizes that genetic diversity of sea ice viruses remains largely unexplored. Similarly, ocean viromes contain high number of sequences having no homologs in reference databases (76).

In addition to just a handful of sea ice virus isolates with sequenced genomes, metagenomics-based studies addressing viral diversity in the Southern Ocean are also still scarce (19, 73). Exploring metagenomes-derived viral sequences deposited in the IMG/VR database with four Antarctic sea ice virus genomes as queries showed that similar sequences may be found across different geographically distant environments. Obviously, ice melting may increase the transmission of viruses from Antarctica, e.g., from ancient glacial lakes to sea ice and seawater. It is, thus, intriguing to see whether future samplings performed in Antarctica and beyond would shed light on the global distribution of viruses related to the Antarctic sea ice phages studied here.

## Acknowledgements

Authors sincerely thank Sari Korhonen, Roselia Henriksson, and Heli Marttila for skillful technical assistance. Dr. Ilona Rissanen is acknowledged for her advice in negative staining. We acknowledge Electron Microscopy Unit (EMBI) and DNA Genomics and Sequencing core facility, Helsinki Institute of Life Science, University of Helsinki. The facilities and expertise of the HiLIFE Biocomplex unit at the University of Helsinki, a member of Instruct-ERIC Centre Finland, FINStruct, and Biocenter Finland are gratefully acknowledged. We thank the Nessling Foundation (T.D.), the Kone Foundation (T.D.), and the Academy of Finland (grant 330977 to T.D.). The work conducted by the U.S. Department of Energy Joint Genome Institute (S.R.) is supported by the Office of Science of the U.S. Department of Energy under contract no. DE-AC02-05CH11231.

## Competing interests

Authors declare no competing financial interests in relation to the work described.

## Notes

### Competing Interest Statement

The authors have declared no competing interest.

### Summary of Updates

Abstract and text edited.

## References

1. Vancoppenolle M, Meiners KM, Michel C, Bopp L, Brabant F, Carnat G, et al. Role of sea ice in global biogeochemical cycles: emerging views and challenges. Quaternary science reviews. 2013;79:207–30.

2. Dieckmann GS, Hellmer HH. The importance of sea ice: an overview. Sea ice. 2010;2:1–22.

3. Arrigo KR. Sea ice ecosystems. Annual review of marine science. 2014;6:439–67.

4. Boetius A, Anesio AM, Deming JW, Mikucki JA, Rapp JZ. Microbial ecology of the cryosphere: sea ice and glacial habitats. Nature Reviews Microbiology. 2015;13(11):677–90.

5. Ewert M, Deming JW. Sea ice microorganisms: Environmental constraints and extracellular responses. Biology. 2013;2(2):603–28.

6. Maccario L, Sanguino L, Vogel TM, Larose C. Snow and ice ecosystems: not so extreme. Research in microbiology. 2015;166(10):782–95.

7. Thomas D, Dieckmann G. Antarctic sea ice--a habitat for extremophiles. Science. 2002;295(5555):641–4.

8. Mock T, Thomas DN. Recent advances in sea‐ice microbiology. Environmental Microbiology. 2005;7(5):605–19.

9. Chénard C, Lauro FM. Exploring the viral ecology of high latitude aquatic systems. Microbial Ecology of Extreme Environments: Springer; 2017. p. 185–200.

10. López-Bueno A, Tamames J, Velázquez D, Moya A, Quesada A, Alcamí A. High diversity of the viral community from an Antarctic lake. Science. 2009;326(5954):858–61.

11. Cavicchioli R. Microbial ecology of Antarctic aquatic systems. Nature Reviews Microbiology. 2015;13(11):691–706.

12. Maranger R, Bird DF, Juniper SK. Viral and bacterial dynamics in Arctic sea ice during the spring algal bloom near Resolute, NWT, Canada. Marine Ecology Progress Series. 1994:121–7.

13. Gowing MM, Riggs BE, Garrison DL, Gibson AH, Jeffries MO. Large viruses in Ross Sea late autumn pack ice habitats. Marine Ecology Progress Series. 2002;241:1–11.

14. Gowing M. Large viruses and infected microeukaryotes in Ross Sea summer pack ice habitats. Marine Biology. 2003;142(5):1029–40.

15. Gowing MM, Garrison DL, Gibson AH, Krupp JM, Jeffries MO, Fritsen CH. Bacterial and viral abundance in Ross Sea summer pack ice communities. Marine Ecology Progress Series. 2004;279:3–12.

16. Marchant H, Davidson A, Wright S, Glazebrook J. The distribution and abundance of viruses in the Southern Ocean during spring. Antarctic Science. 2000;12(4):414–7.

17. Wells LE, Deming JW. Modelled and measured dynamics of viruses in Arctic winter sea‐ice brines. Environmental Microbiology. 2006;8(6):1115–21.

18. Säwström C, Lisle J, Anesio AM, Priscu JC, Laybourn-Parry J. Bacteriophage in polar inland waters. Extremophiles. 2008;12(2):167–75.

19. Brum JR, Hurwitz BL, Schofield O, Ducklow HW, Sullivan MB. Seasonal time bombs: dominant temperate viruses affect Southern Ocean microbial dynamics. The ISME journal. 2016;10(2):437–49.

20. Anesio AM, Bellas CM. Are low temperature habitats hot spots of microbial evolution driven by viruses? Trends in microbiology. 2011;19(2):52–7.

21. Yu Z-C, Chen X-L, Shen Q-T, Zhao D-L, Tang B-L, Su H-N, et al. Filamentous phages prevalent in Pseudoalteromonas spp. confer properties advantageous to host survival in Arctic sea ice. The ISME journal. 2015;9(4):871–81.

22. Borriss M, Helmke E, Hanschke R, Schweder T. Isolation and characterization of marine psychrophilic phage-host systems from Arctic sea ice. Extremophiles. 2003;7(5):377–84.

23. Luhtanen A-M, Eronen-Rasimus E, Kaartokallio H, Rintala J-M, Autio R, Roine E. Isolation and characterization of phage–host systems from the Baltic Sea ice. Extremophiles. 2014;18(1):121–30.

24. Luhtanen A-M, Eronen-Rasimus E, Oksanen HM, Tison J-L, Delille B, Dieckmann GS, et al. The first known virus isolates from Antarctic sea ice have complex infection patterns. FEMS microbiology ecology. 2018;94(4):fiy028.

25. Senčilo A, Luhtanen AM, Saarijärvi M, Bamford DH, Roine E. Cold‐active bacteriophages from the Baltic Sea ice have diverse genomes and virus–host interactions. Environmental microbiology. 2015;17(10):3628–41.

26. Tedesco L, Vichi M. Sea ice biogeochemistry: A guide for modellers. PloS one. 2014;9(2):e89217.

27. Adams MH. Bacteriophages: Citeseer; 1959.

28. Schneider CA, Rasband WS, Eliceiri KW. NIH Image to ImageJ: 25 years of image analysis. Nature methods. 2012;9(7):671–5.

29. Bankevich A, Nurk S, Antipov D, Gurevich AA, Dvorkin M, Kulikov AS, et al. SPAdes: a new genome assembly algorithm and its applications to single-cell sequencing. Journal of computational biology. 2012;19(5):455–77.

30. Madeira F, Park YM, Lee J, Buso N, Gur T, Madhusoodanan N, et al. The EMBL-EBI search and sequence analysis tools APIs in 2019. Nucleic acids research. 2019;47(W1):W636–W41.

31. Altschul SF, Gish W, Miller W, Myers EW, Lipman DJ. Basic local alignment search tool. Journal of molecular biology. 1990;215(3):403–10.

32. Lu S, Wang J, Chitsaz F, Derbyshire MK, Geer RC, Gonzales NR, et al. CDD/SPARCLE: the conserved domain database in 2020. Nucleic acids research. 2020;48(D1):D265–D8.

33. Zimmermann L, Stephens A, Nam S-Z, Rau D, Kübler J, Lozajic M, et al. A completely reimplemented MPI bioinformatics toolkit with a new HHpred server at its core. Journal of molecular biology. 2018;430(15):2237–43.

34. Lopes A, Tavares P, Petit M-A, Guérois R, Zinn-Justin S. Automated classification of tailed bacteriophages according to their neck organization. BMC genomics. 2014;15(1):1–17.

35. Chan PP, Lowe TM. tRNAscan-SE: searching for tRNA genes in genomic sequences. Gene Prediction: Springer; 2019. p. 1–14.

36. Roux S, Páez-Espino D, Chen I-MA, Palaniappan K, Ratner A, Chu K, et al. IMG/VR v3: an integrated ecological and evolutionary framework for interrogating genomes of uncultivated viruses. Nucleic Acids Research. 2021;49(D1):D764–D75.

37. Darzentas N. Circoletto: visualizing sequence similarity with Circos. Bioinformatics. 2010;26(20).

38. Sullivan MJ, Petty NK, Beatson SA. Easyfig: a genome comparison visualizer. Bioinformatics. 2011;27(7):1009–10.

39. Christiam C, George C, Vahram A, Ning M, Jason P, Kevin B, et al. BLAST+: architecture and applications. BMC bioinformatics. 2009;10(1):421.

40. Adriaenssens EM, Cowan DA. Using signature genes as tools to assess environmental viral ecology and diversity. Applied and environmental microbiology. 2014;80(15):4470–80.

41. Henthorn KS, Friedman DI. Identification of related genes in phages phi 80 and P22 whose products are inhibitory for phage growth in Escherichia coli IHF mutants. Journal of bacteriology. 1995;177(11):3185–90.

42. Casjens SR, Gilcrease EB, Winn-Stapley DA, Schicklmaier P, Schmieger H, Pedulla ML, et al. The generalized transducing Salmonella bacteriophage ES18: complete genome sequence and DNA packaging strategy. Journal of bacteriology. 2005;187(3):1091–104.

43. Szewczyk B, Bienkowska-Szewczyk K, Kozloff LM. Identification of T4 gene 25 product, a component of the tail baseplate, as a 15K lysozyme. Molecular and General Genetics MGG. 1986;202(3):363–7.

44. Petrov VM, Ratnayaka S, Nolan JM, Miller ES, Karam JD. Genomes of the T4-related bacteriophages as windows on microbial genome evolution. Virology journal. 2010;7(1):1–19.

45. Kavagutti VS, Andrei A-Ş, Mehrshad M, Salcher MM, Ghai R. Phage-centric ecological interactions in aquatic ecosystems revealed through ultra-deep metagenomics. Microbiome. 2019;7(1):1–15.

46. Panwar P, Allen MA, Williams TJ, Hancock AM, Brazendale S, Bevington J, et al. Influence of the polar light cycle on seasonal dynamics of an Antarctic lake microbial community. Microbiome. 2020;8(1):1–24.

47. Hawley ER, Malfatti SA, Pagani I, Huntemann M, Chen A, Foster B, et al. Metagenomes from two microbial consortia associated with Santa Barbara seep oil. Marine genomics. 2014;18:97–9.

48. Hawley ER, Piao H, Scott NM, Malfatti S, Pagani I, Huntemann M, et al. Metagenomic analysis of microbial consortium from natural crude oil that seeps into the marine ecosystem offshore Southern California. Standards in Genomic Sciences. 2014;9(3):1259–74.

49. Sun M, Zhan Y, Marsan D, Páez-Espino D, Cai L, Chen F. Uncultivated Viral Populations Dominate Estuarine Viromes on the Spatiotemporal Scale. Msystems. 2021;6(2):e01020–20.

50. Tschitschko B, Erdmann S, DeMaere MZ, Roux S, Panwar P, Allen MA, et al. Genomic variation and biogeography of Antarctic haloarchaea. Microbiome. 2018;6(1):1–16.

51. Williams TJ, Allen MA, Ivanova N, Huntemann M, Haque S, Hancock AM, et al. Genome Analysis of a Verrucomicrobial Endosymbiont With a Tiny Genome Discovered in an Antarctic Lake. Frontiers in microbiology. 2021;12.

52. Ji X, Yu H, Zhang Q, Lin L, Wei Y. Isolation and characterization of a novel lytic cold-active bacteriophage VNPH-1 from the Napahai wetland in China. Annals of microbiology. 2015;65(3):1789–96.

53. Ji X, Zhang C, Fang Y, Zhang Q, Lin L, Tang B, et al. Isolation and characterization of glacier VMY22, a novel lytic cold-active bacteriophage of Bacillus cereus. Virologica Sinica. 2015;30(1):52–8.

54. Li M, Wang J, Zhang Q, Lin L, Kuang A, Materon LA, et al. Isolation and characterization of the lytic cold-active bacteriophage MYSP06 from the Mingyong glacier in China. Current microbiology. 2016;72(2):120–7.

55. Qin K, Ji X, Zhang C, Ding Y, Kuang A, Zhang S, et al. Isolation and characterization of wetland VSW-3, a novel lytic cold-active bacteriophage of Pseudomonas fluorescens. Canadian journal of microbiology. 2017;63(2):110–8.

56. Xiang Y, Wang S, Li J, Wei Y, Zhang Q, Lin L, et al. Isolation and characterization of two lytic cold-active bacteriophages infecting Pseudomonas fluorescens from the Napahai plateau wetland. Canadian journal of microbiology. 2018;64(3):183–90.

57. Wells LE, Deming JW. Effects of temperature, salinity and clay particles on inactivation and decay of cold-active marine Bacteriophage 9A. Aquatic microbial ecology. 2006;45(1):31–9.

58. Kukkaro P, Bamford DH. Virus–host interactions in environments with a wide range of ionic strengths. Environmental microbiology reports. 2009;1(1):71–7.

59. Rodela ML, Sabet S, Peterson A, Dillon JG. Broad environmental tolerance for a Salicola host-phage pair isolated from the Cargill Solar Saltworks, Newark, CA, USA. Microorganisms. 2019;7(4):106.

60. Aalto AP, Bitto D, Ravantti JJ, Bamford DH, Huiskonen JT, Oksanen HM. Snapshot of virus evolution in hypersaline environments from the characterization of a membrane-containing Salisaeta icosahedral phage 1. Proceedings of the National Academy of Sciences. 2012;109(18):7079–84.

61. Jończyk E, Kłak M, Międzybrodzki R, Górski A. The influence of external factors on bacteriophages. Folia microbiologica. 2011;56(3):191–200.

62. Vörös Z, Csík G, Herényi L, Kellermayer M. Temperature-dependent nanomechanics and topography of bacteriophage T7. Journal of virology. 2018;92(20).

63. Langlet J, Gaboriaud F, Gantzer C. Effects of pH on plaque forming unit counts and aggregation of MS2 bacteriophage. Journal of Applied Microbiology. 2007;103(5):1632–8.

64. Szermer-Olearnik B, Drab M, Mąkosa M, Zembala M, Barbasz J, Dąbrowska K, et al. Aggregation/dispersion transitions of T4 phage triggered by environmental ion availability. Journal of nanobiotechnology. 2017;15(1):1–15.

65. Liu Y, Wang J, Liu Y, Wang Y, Zhang Z, Oksanen HM, et al. Identification and characterization of SNJ 2, the first temperate pleolipovirus integrating into the genome of the SNJ 1‐lysogenic archaeal strain. Molecular microbiology. 2015;98(6):1002–20.

66. Laybourn-Parry J, Marshall WA, Madan NJ. Viral dynamics and patterns of lysogeny in saline Antarctic lakes. Polar Biology. 2007;30(3):351–8.

67. Säwström C, Anesio MA, Granéli W, Laybourn-Parry J. Seasonal viral loop dynamics in two large ultraoligotrophic Antarctic freshwater lakes. Microbial ecology. 2007;53(1):1–11.

68. Mäntynen S, Laanto E, Oksanen HM, Poranen MM, Díaz-Muñoz SL. Black box of phage–bacterium interactions: exploring alternative phage infection strategies. Open biology. 2021;11(9):210188.

69. Middelboe M, Nielsen TG, Bjørnsen PK. Viral and bacterial production in the North Water: in situ measurements, batch-culture experiments and characterization and distribution of a virus–host system. Deep Sea Research Part II: Topical Studies in Oceanography. 2002;49(22–23):5063–79.

70. Delisle A, Levin R. Effect of temperature on an obligately psychrophilic phage-host system of Pseudomonas putrefaciens. Antonie Van Leeuwenhoek. 1972;38(1):9–15.

71. Wells LE, Deming JW. Characterization of a cold-active bacteriophage on two psychrophilic marine hosts. Aquatic microbial ecology. 2006;45(1):15–29.

72. Sillankorva S, Oliveira R, Vieira MJ, Sutherland I, Azeredo J. Pseudomonas fluorescens infection by bacteriophage ΦS1: the influence of temperature, host growth phase and media. FEMS microbiology letters. 2004;241(1):13–20.

73. Brum JR, Ignacio-Espinoza JC, Roux S, Doulcier G, Acinas SG, Alberti A, et al. Patterns and ecological drivers of ocean viral communities. Science. 2015;348(6237).

74. Colangelo-Lillis JR, Deming JW. Genomic analysis of cold-active Colwelliaphage 9A and psychrophilic phage–host interactions. Extremophiles. 2013;17(1):99–114.

75. Brum JR, Sullivan MB. Rising to the challenge: accelerated pace of discovery transforms marine virology. Nature Reviews Microbiology. 2015;13(3):147–59.

76. Hurwitz BL, Sullivan MB. The Pacific Ocean Virome (POV): a marine viral metagenomic dataset and associated protein clusters for quantitative viral ecology. PloS one. 2013;8(2):e57355.

